## Supplemental tables and figures for "Cannabinoid Combination Targets *NOTCH1*-Mutated T-ALL Through the Integrated Stress Response Pathway"

**This PDF file includes:**

table S1. Phytocannabinoid concentrations by UHPLC/LC-MS of fractions relative to the whole extract.

table S2. Phytocannabinoid concentrations by UHPLC/LC-MS of cannabis fraction 2 peaks.

table S3. $^{1}H$ and $^{13}C$ peak assignments and chemical shifts of 331-18A.

table S4. Ten most increased- and decreased-abundance genes following treatment of MOLT-4 cells with the whole extract according to Affymetrix.

fig. S1. Fraction 2 is as effective as the whole extract in reducing the viability of MOLT-4 cells.

fig. S2. Specific identification of phytocannabinoids with anti-cancer properties by spectral matching and peak purity.

fig. S3. CBD and 331-18A NMR spectra.

fig. S4. Notch1 downregulation by Extract 12 is mediated through CB2 and TRPV1 followed by ATF4-CHOP-CHAC1 signaling pathway.

fig. S5. Whole extract treatment activates eIF2α.

fig. S6. Inhibition of tumor growth *in-vivo* by whole extract.

Fig. S7. Treatment with extract containing the three cannabinoids inhibits leukemic expansion in an *in-vivo* PDX model.

**Supplementary table S1. Phytocannabinoid concentrations by UHPLC/LC-MS of fractions relative to the whole extract**

| | | Fraction | | | |
|---|---|---|---|---|---|
| [% w/w] | Whole extract | 1 | 2 | 3 | 4 |
| **CBC** | 2.856 | 0.000 | 0.000 | 32.607 | 0.099 |
| **CBCA** | 0.012 | 0.000 | 0.000 | 0.001 | 0.153 |
| **CBCA-C4** | 0.000 | 0.000 | 0.000 | 0.001 | 0.000 |
| **CBC-C4** | 0.023 | 0.000 | 0.000 | 0.248 | 0.000 |
| **CBCMA** | 0.004 | 0.000 | 0.003 | 0.004 | 0.000 |
| **CBCO** | 0.002 | 0.000 | 0.003 | 0.000 | 0.000 |
| **CBCOA** | 0.000 | 0.000 | 0.000 | 0.000 | 0.000 |
| **CBCV** | 0.267 | 0.000 | 0.000 | 3.028 | 0.000 |
| **CBCVA** | 0.002 | 0.000 | 0.000 | 0.008 | 0.000 |
| **CBD** | 53.384 | 0.018 | 77.534 | 0.045 | 0.034 |
| **CBDA** | 1.338 | 0.059 | 1.800 | 0.187 | 0.298 |
| **CBDA-C4** | 0.006 | 0.001 | 0.009 | 0.002 | 0.001 |
| **CBD-C4** | 0.256 | 0.000 | 0.396 | 0.000 | 0.000 |
| **CBDM** | 0.014 | 0.000 | 0.005 | 0.138 | 0.000 |
| **CBDMA** | 0.000 | 0.000 | 0.000 | 0.000 | 0.000 |
| **CBDO** | 0.004 | 0.004 | 0.007 | 0.000 | 0.000 |
| **CBDOA** | 0.000 | 0.001 | 0.001 | 0.000 | 0.000 |
| **CBDV** | 2.929 | 0.000 | 4.416 | 0.000 | 0.000 |
| **CBDVA** | 0.055 | 0.003 | 0.084 | 0.013 | 0.006 |
| **CBE** | 0.339 | 0.050 | 0.544 | 0.000 | 0.000 |
| **CBEA** | 0.010 | 0.001 | 0.014 | 0.002 | 0.002 |
| **CBEV** | 0.014 | 0.005 | 0.020 | 0.000 | 0.000 |
| **CBEVA** | 0.001 | 0.003 | 0.001 | 0.000 | 0.000 |
| **CBG** | 0.990 | 0.000 | 1.544 | 0.000 | 0.000 |
| **CBGA** | 0.024 | 0.000 | 0.032 | 0.004 | 0.007 |
| **CBGA-C4** | 0.000 | 0.000 | 0.000 | 0.000 | 0.000 |
| **CBG-C4** | 0.002 | 0.000 | 0.002 | 0.000 | 0.000 |
| **CBGM** | 0.001 | 0.000 | 0.002 | 0.000 | 0.000 |
| **CBGMA** | 0.000 | 0.000 | 0.000 | 0.000 | 0.000 |
| **CBGO** | 0.000 | 0.000 | 0.000 | 0.000 | 0.000 |
| **CBGOA** | 0.000 | 0.001 | 0.000 | 0.000 | 0.000 |
| **CBGV** | 0.008 | 0.000 | 0.013 | 0.000 | 0.000 |
| **CBGVA** | 0.000 | 0.000 | 0.000 | 0.000 | 0.000 |
| **CBL** | 0.026 | 0.000 | 0.000 | 0.327 | 0.000 |

| | | | | | |
|---|---|---|---|---|---|
| **CBN** | 0.118 | 0.013 | 0.002 | 1.479 | 0.002 |
| **CBNA** | 0.000 | 0.000 | 0.000 | 0.000 | 0.000 |
| **CBNA-C4** | 0.000 | 0.000 | 0.000 | 0.000 | 0.000 |
| **CBN-C4** | 0.000 | 0.000 | 0.001 | 0.000 | 0.000 |
| **CBND** | 0.008 | 0.000 | 0.015 | 0.000 | 0.000 |
| **CBNDA** | 0.001 | 0.000 | 0.001 | 0.000 | 0.000 |
| **CBNDVA** | 0.000 | 0.001 | 0.000 | 0.000 | 0.000 |
| **CBNM** | 0.000 | 0.000 | 0.000 | 0.000 | 0.000 |
| **CBNMA** | 0.000 | 0.000 | 0.000 | 0.000 | 0.000 |
| **CBNO** | 0.000 | 0.000 | 0.000 | 0.000 | 0.000 |
| **CBNOA** | 0.000 | 0.000 | 0.000 | 0.000 | 0.000 |
| **CBNV** | 0.005 | 0.000 | 0.009 | 0.000 | 0.000 |
| **CBNVA** | 0.000 | 0.000 | 0.000 | 0.000 | 0.000 |
| **CBT-1** | 0.065 | 0.016 | 0.099 | 0.086 | 0.008 |
| **CBT-2** | 0.015 | 0.004 | 0.015 | 0.028 | 0.001 |
| **CBT-3** | 0.047 | 0.023 | 0.067 | 0.128 | 0.009 |
| **CBTA-1** | 0.000 | 0.001 | 0.000 | 0.000 | 0.000 |
| **CBTA-3** | 0.000 | 0.000 | 0.002 | 0.001 | 0.000 |
| **CBTV-1** | 0.004 | 0.139 | 0.000 | 0.000 | 0.000 |
| **CBTV-3** | 0.000 | 0.013 | 0.002 | 0.000 | 0.000 |
| **d8-THC** | 0.014 | 0.000 | 0.000 | 0.159 | 0.000 |
| **d9-THC** | 1.620 | 0.000 | 0.000 | 19.001 | 0.000 |
| **d9-THCA** | 0.000 | 0.000 | 0.000 | 0.000 | 0.000 |
| **d9-THCA-C4** | 0.000 | 0.000 | 0.000 | 0.000 | 0.000 |
| **d9-THC-C4** | 0.000 | 0.000 | 0.000 | 0.084 | 0.000 |
| **d9-THCM** | 0.000 | 0.000 | 0.000 | 0.000 | 0.000 |
| **d9-THCMA** | 0.000 | 0.000 | 0.000 | 0.000 | 0.000 |
| **d9-THCO** | 0.000 | 0.000 | 0.000 | 0.000 | 0.000 |
| **d9-THCOA** | 0.000 | 0.000 | 0.000 | 0.000 | 0.000 |
| **d9-THCV** | 0.150 | 0.000 | 0.245 | 0.000 | 0.000 |
| **d9-THCVA** | 0.000 | 0.000 | 0.000 | 0.000 | 0.000 |
| **OH-CBN** | 0.003 | 0.000 | 0.004 | 0.015 | 0.000 |
| **OH-CBNA** | 0.000 | 0.000 | 0.000 | 0.000 | 0.000 |
| **SesquiCBG** | 0.025 | 0.000 | 0.000 | 0.228 | 0.000 |
| **SesquiCBGA** | 0.000 | 0.000 | 0.000 | 0.000 | 0.000 |
| **313-16b** | 0.273 | 0.000 | 0.000 | 2.981 | 0.000 |
| **327-13a** | 0.201 | 0.006 | 0.330 | 0.015 | 0.000 |
| **327-13b** | 0.178 | 0.012 | 0.270 | 0.023 | 0.000 |
| **327-13c** | 0.249 | 0.064 | 0.174 | 0.830 | 0.047 |
| **329-11a** | 0.011 | 0.592 | 0.001 | 0.000 | 0.000 |
| **329-11b** | 0.118 | 0.002 | 0.194 | 0.000 | 0.000 |
| **329-11c** | 0.016 | 0.007 | 0.020 | 0.170 | 0.000 |

| | | | | | |
|---|---|---|---|---|---|
| **329-11d** | 0.000 | 0.000 | 0.000 | 0.002 | 0.000 |
| **331-18a** | 2.042 | 0.003 | 3.308 | 0.003 | 0.000 |
| **331-18b** | 0.543 | 0.000 | 0.511 | 0.007 | 0.037 |
| **331-18c** | 0.083 | 0.130 | 0.126 | 0.029 | 0.098 |
| **331-18d** | 0.073 | 0.000 | 0.010 | 0.436 | 0.000 |
| **357-16a** | 0.000 | 0.000 | 0.000 | 0.000 | 0.000 |
| **361-17a** | 0.003 | 0.019 | 0.005 | 0.002 | 0.000 |
| **361-17b** | 0.003 | 0.125 | 0.000 | 0.002 | 0.000 |
| **371-14a** | 0.002 | 0.002 | 0.003 | 0.001 | 0.001 |
| **371-14b** | 0.000 | 0.000 | 0.000 | 0.000 | 0.000 |
| **373-12a** | 0.000 | 0.004 | 0.000 | 0.000 | 0.000 |
| **373-12b** | 0.000 | 0.000 | 0.000 | 0.000 | 0.001 |
| **373-12c** | 0.000 | 0.000 | 0.000 | 0.000 | 0.000 |
| **373-12d** | 0.000 | 0.000 | 0.000 | 0.000 | 0.000 |
| **373-15b** | 0.000 | 0.002 | 0.000 | 0.003 | 0.000 |
| **373-15c** | 0.383 | 0.013 | 0.002 | 0.021 | 4.478 |
| **375-19a** | 0.003 | 0.016 | 0.004 | 0.002 | 0.001 |
| **375-19b** | 0.000 | 0.005 | 0.000 | 0.000 | 0.000 |
| **375-19c** | 0.000 | 0.000 | 0.000 | 0.000 | 0.000 |
| **417-15a** | 0.000 | 0.001 | 0.000 | 0.000 | 0.000 |

**Supplementary table S2. Phytocannabinoid concentrations by UHPLC/LC-MS of cannabis fraction 2 peaks**

| | P1 | P2 | P3 | P4 | P5 |
|---|---|---|---|---|---|
| **331-18a** | 181.0502 | 5.0040 | 0.1243 | 0.1195 | 0.0000 |
| **CBD** | 4.0371 | 0.0915 | 1.0801 | 1.0917 | 109.3740 |
| **CBDVA** | 1.1604 | 0.0411 | 0.0000 | 0.0000 | 0.0000 |
| **CBDA** | 0.9107 | 0.0000 | 0.0385 | 11.7560 | 0.0000 |
| **CBND** | 0.6878 | 0.0307 | 0.0000 | 0.0000 | 0.0000 |
| **THCA** | 0.5969 | 0.0000 | 0.1629 | 0.1648 | 0.0000 |
| **329-11b** | 0.3289 | 0.2398 | 0.0089 | 0.0151 | 0.0000 |
| **CBT-2** | 0.1802 | 0.0135 | 0.0152 | 0.0147 | 0.0000 |
| **373-15c** | 0.1545 | 0.0876 | 0.0558 | 0.0565 | 0.0000 |
| **327-13c** | 0.0546 | 0.0000 | 0.0192 | 0.0493 | 0.1371 |
| **CBCA** | 0.0401 | 0.0000 | 0.0000 | 0.0000 | 0.0000 |
| **CBDV** | 0.0246 | 93.2161 | 0.8774 | 0.9112 | 0.0000 |
| **327-13b** | 0.0212 | 0.0000 | 0.1556 | 0.1526 | 0.0000 |
| **CBN** | 0.0152 | 0.0000 | 0.0989 | 0.0870 | 0.0000 |
| **327-13a** | 0.0000 | 0.9210 | 0.1657 | 0.1922 | 0.0000 |
| **CBGV** | 0.0000 | 0.2160 | 0.0000 | 0.0000 | 0.0000 |
| **CBD-C4** | 0.0000 | 0.0000 | 20.5724 | 0.1509 | 0.0000 |
| **CBT-3** | 0.0000 | 0.0000 | 0.0783 | 0.0844 | 0.0000 |
| **361-17a** | 0.0000 | 0.0000 | 0.0438 | 0.0416 | 0.0000 |
| **CBG-C4** | 0.0000 | 0.0000 | 0.0254 | 0.0262 | 0.0000 |
| **373-15b** | 0.0000 | 0.0000 | 0.0244 | 0.0231 | 0.0000 |
| **329-11d** | 0.0000 | 0.0000 | 0.0179 | 0.0198 | 0.0000 |
| **CBEA** | 0.0000 | 0.0000 | 0.0135 | 0.0361 | 0.0000 |
| **CBGA-C4** | 0.0000 | 0.0000 | 0.0079 | 0.0079 | 0.0000 |
| **CBDA-C4** | 0.0000 | 0.0000 | 0.0030 | 0.0030 | 0.0000 |
| **331-18b** | 0.0000 | 0.0000 | 0.0000 | 2.0747 | 0.3356 |
| **CBNV** | 0.0000 | 0.0000 | 0.0000 | 0.0338 | 0.0000 |
| **CBG** | 0.0000 | 0.0000 | 0.0000 | 0.0000 | 0.4119 |
| **CBE** | 0.0000 | 0.0000 | 0.0000 | 0.0000 | 0.0142 |

**Supplementary table S3: $^{1}$H and $^{13}$C peak assignments and chemical shifts of CBD and 331-18A**

| | **CBD**[a] | | **331-18A** | |
|---|---|---|---|---|
| | **$^{1}$H** | **$^{13}$C** | **$^{1}$H** | **$^{13}$C** |
| **1** | 3.85 | 37.08 | 3.83 | 32.63 |
| **2** | 5.57 | 123.96 | 5.71 | 123.45 |
| **3** | - | 140.21 | - | 140.20 |
| **4** | 2.10, 2.23 | 30.31 | 2.06, 2.13 | 27.08 |
| **5** | 1.77, 1.82 | 28.28 | 1.73, 1.97 | 22.66 |
| **6** | 2.39 | 46.10 | 1.90 | 48.24 |
| **7** | 1.79 | 23.75 | 1.81 | 23.82 |
| **8** | - | 149.38 | - | 75.19 |
| **9** | 1.66 | 20.44 | 1.25 | 25.97 |
| **10** | 4.55, 4.67 | 110.86 | 1.26 | 29.70 |
| **1'** | - | 113.63 | - | 114.66 |
| **2'** | - | 155.95 | - | 155.97 |
| **2'-OH** | 6.05 | - | 6.61 | - |
| **3'** | 6.29 | 109.68 | 6.26 | 109.44 |
| **4'** | - | 143.02 | - | 143.52 |
| **5'** | 6.16 | 107.88 | 6.33 | 109.44 |
| **5'-OH** | 4.77 | - | 7.64 | - |
| **6'** | - | 153.78 | - | 154.23 |
| **1''** | 2.43 | 35.44 | 2.45 | 35.49 |
| **2''** | 1.55 | 30.69 | 1.57 | 30.74 |
| **3''** | 1.27 | 31.46 | 1.29 | 31.52 |
| **4''** | 1.30 | 22.55 | 1.31 | 22.56 |
| **5''** | 0.87 | 14.09 | 0.88 | 14.08 |

[a] Peak assignments are in close agreement with the literature (*33*).

**Supplementary table S4. Ten most increased- and decreased-abundance genes following treatment of MOLT-4 cells with the whole extract according to Affymetrix**

| | Increased abundance [a] | | Decreased abundance [a] | |
|---|---|---|---|---|
| **1** | *SLC7A11* | 34.2 | *EYA4* | -7.25 |
| **2** | *CHAC1* | 27.48 | *CHTF8* | -6.83 |
| **3** | *JUN* | 25.75 | *CD180* | -5.74 |
| **4** | *ID2* | 21.51 | *DHCR7* | -5.51 |
| **5** | *MT1L* | 19.88 | *FSIP1* | -5.28 |
| **6** | *SNAI1* | 17.41 | *DEFB113* | -5.25 |
| **7** | *SLC43A1* | 16.96 | *C15orf65* | -5.22 |
| **8** | *MT1X* | 16.57 | *KRTAP10-10* | -5.2 |
| **9** | *EGR1* | 16.03 | *GRIA2* | -5.11 |
| **10** | *CITED2* | 15.98 | *OR5T3* | -5.02 |

[a] Results are presented as gene expression fold change compared to vehicle treatment. The full list of genes that are up-regulated or down-regulated upon treatment with Extract 12 is available in the Gene Expression Omnibus (GEO) repository, GSE154287, https://www.ncbi.nlm.nih.gov/geo/query/acc.cgi?acc=GSE154287.

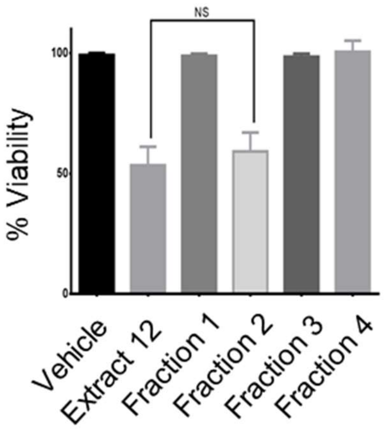


**fig. S1. Fraction 2 is as effective as the whole extract in reducing the viability of MOLT-4 cells**. MOLT-4 cells were treated with either whole extract or each of the four fractions and 24 hrs later the viability of the cells was assessed with XTT.

A

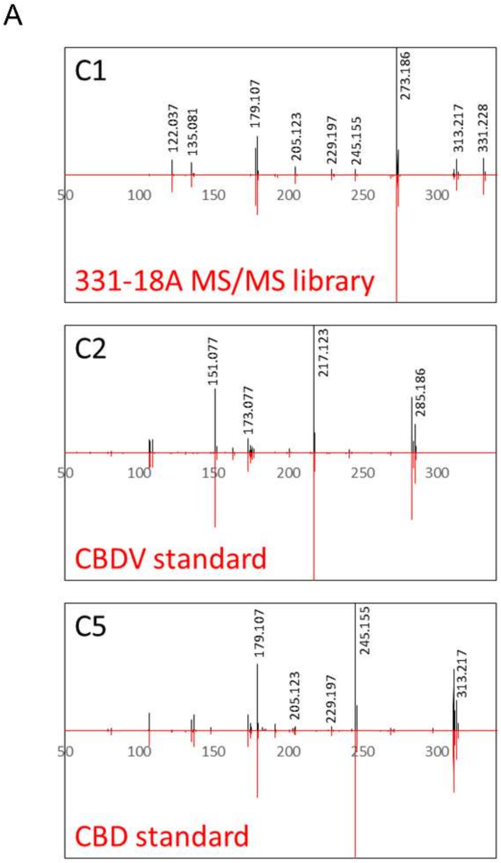


B

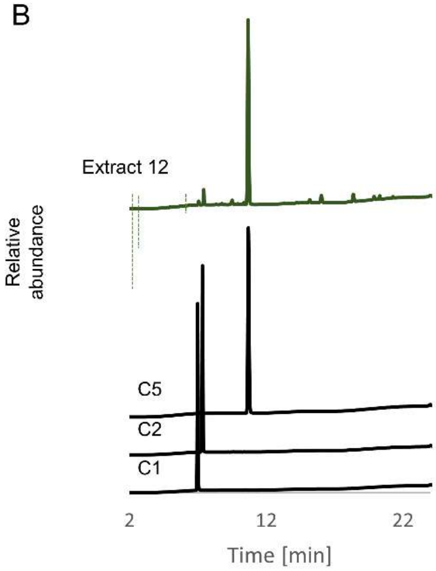


C

| | Identified compound | Absolute purity [% w/w] | Relative peak purity [%] |
|---|---|---|---|
| C1 | 331-18A | 86 | 99 |
| C2 | CBDV | 103 | 100 |
| C5 | CBD | 93 | 100 |

**fig. S2. Specific identification of phytocannabinoids by spectral matching and peak purity.** (**A**) MS/MS spectral matching of C1, C2 and C5 versus our spectral library and analytical standards of CBDV and CBD, respectively. (**B**) UHPLC/UV chromatogram of Extract 12 and the three isolated phytocannabinoids analyzed separately. (**C**) Calculated purities of the three isolated phytocannabinoids according to analytical standards (absolute purity) or as the percent area in relation to all other observed peaks in the UHPLC analysis (relative peak purity). The absolute purity of 331-18A was quantified according to the calibration curve of the CBD standard.

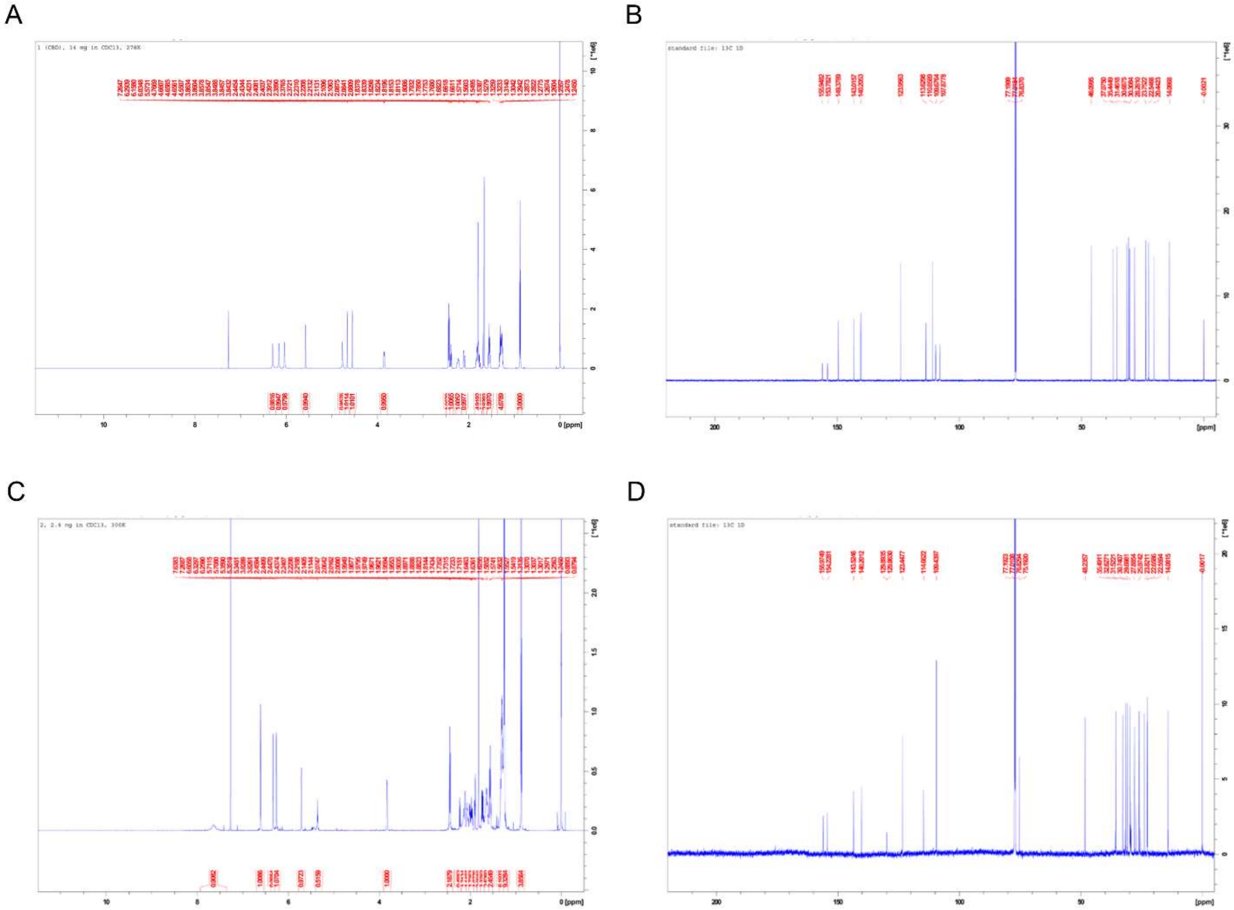


**fig. S3. CBD and 331-18A NMR spectra.** (**A**) $^{1}H$ NMR spectrum of CBD**.** (**B**) $^{13}C$ NMR spectrum of CBD. (**C**) $^{1}H$ NMR spectrum of 331-18A. (**D**) $^{13}C$ NMR spectrum of 331-18A.

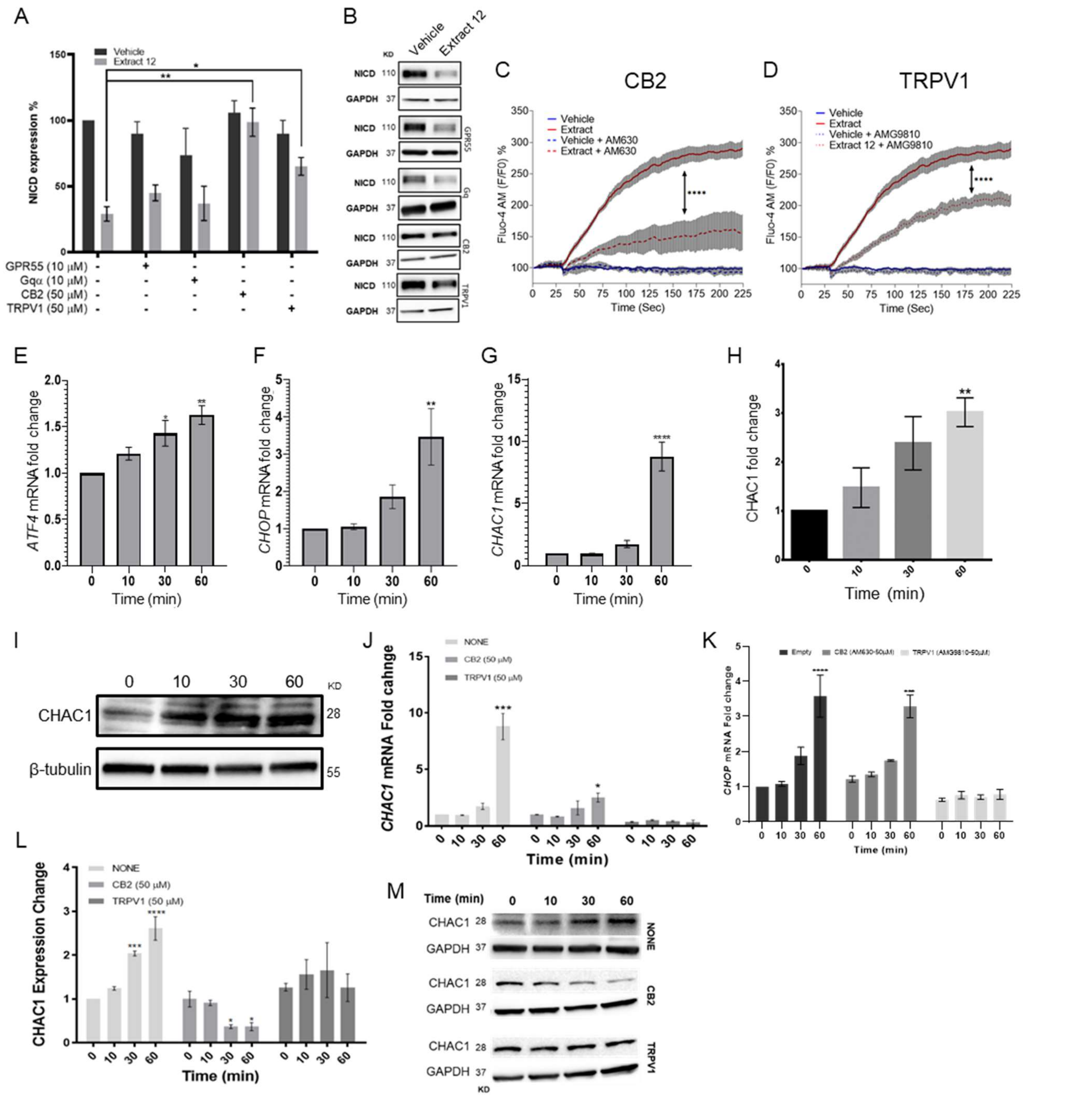


**fig. S4. Notch1 downregulation by Extract 12 is mediated through CB2 and TRPV1 followed by ATF4-CHOP-CHAC1 signaling pathway.** (**A**) MOLT-4 cells were pretreated for 30 minutes with either of the following antagonists: CID (10 µM) for GPR55, BIM (10 µM) for general Gαq GPCR, AM630 (50 µM) for CB2 and AMG9810 (50 µM) for TRPV1; then added vehicle or Extract 12 (3 µg/mL) for 3 hrs. The protein expression of NICD was analyzed relative to untreated vehicle control (n=3) and statistically analyzed with unpaired student's t-test (*p<0.05, **p<0.01). (**B**) Representative blots of NICD expression. (**C-D**) MOLT-4 cells were pretreated for 30 minutes with either AM630 (50 µM), an antagonist to CB2, or AMG9810 (50 µM) an antagonist to TRPV1; then added either vehicle or Extract 12 and calcium release was immediately measured with the Fluo-4 calcium probe. The presented calcium curves represent an average of three independent experiments and statistically analyzed with two-way ANOVA (****p<0.0001). (**E-G**) Timecourse (n=3) qRT-PCR of *ATF4*, *CHOP* and *CHAC1* genes at different time points (0-60 min) following treatment with Extract 12 (3 μg/mL) and statistically analyzed with one-way ANOVA (*p<0.05, **p<0.01, ****p<0.0001). (**H-I**) MOLT-4 cells were treated with Extract 12 (3 μg/mL) for 10, 30 and 60 min and the protein expression of CHAC1 was evaluated with β-tubulin as the loading control. Intensity analysis was performed on three independent experiments relative to untreated control (n=3) and statistically analyzed with an unpaired student's t-test (**p<0.01). (**J-M**) MOLT-4 cells were pretreated for 30 min with vehicle or 50 μM of either antagonist to CB2 (AM630) or to TRPV1 (AMG9810), and then treated with Extract 12 (3 μg/mL) for the indicated times and assessed (n=3) for the mRNA expression of *CHAC1* (J) and *CHOP* (K), as well as the protein expression of CHAC1 (L). A representative blot of CHAC1 expression with GAPDH as the loading control (M). Results are presented as mean ± SEM relative to untreated control and statistically analyzed by unpaired student's t-test (*p<0.05, ***p<0.001, ****p<0.0001).

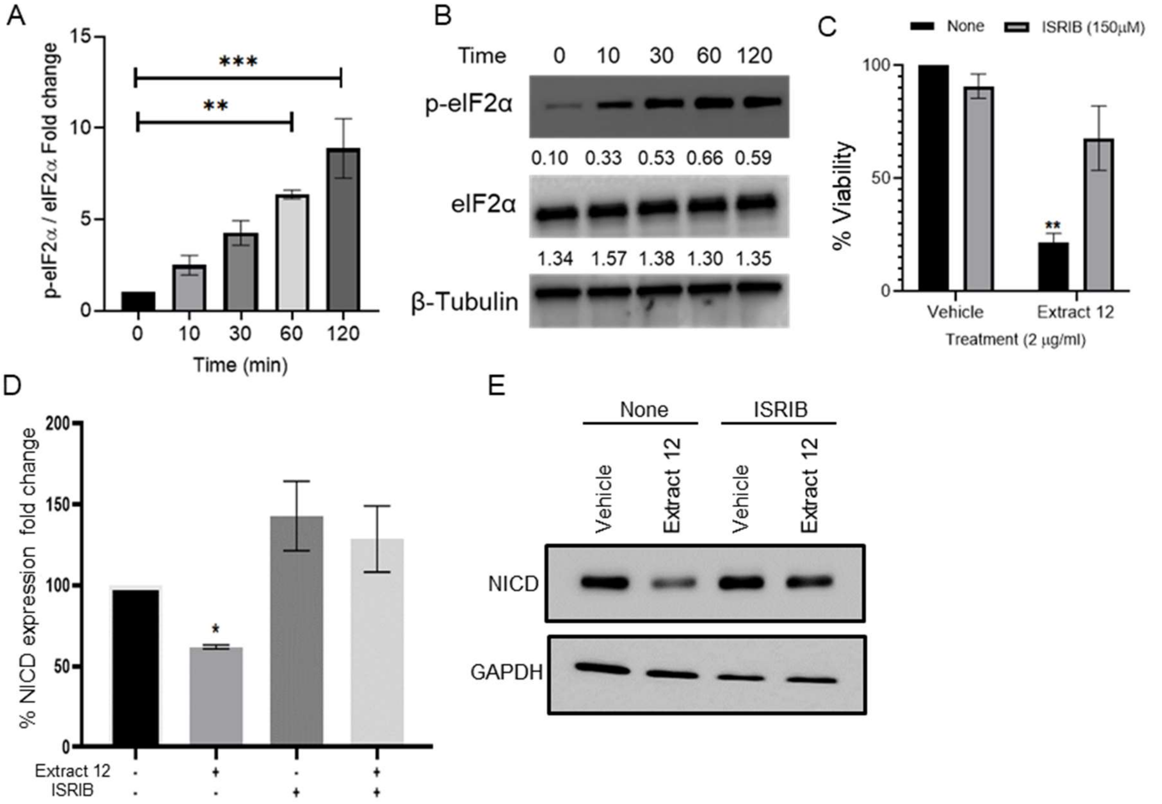


**fig. S5. Whole extract treatment activates eIF2α.** (**A**) MOLT-4 cells were with Extract 12 and the phosphorylation of eIF2α was assessed at different times up to 2 hrs after treatment. (**B**) Representative blots of phosphorylated eIF2α (Ser$^{51}$), total eIF2α and β-Tubulin. (**C**) MOLT-4 cells were pretreated for 30 min with the eIF2α inhibitor ISRIB (150 μM) or left untreated, then treated with vehicle or Extract12. Viability was assessed after 24 hrs with XTT assay (n=3). (**D**) NICD protein expression was assessed (n=3) with GAPDH as the loading control. (**E**) A representative image showing the protein levels of NICD and GAPDH. Results are presented as mean ± SEM and statistically analyzed with one-way ANOVA (*p<0.05, **p<0.01, ***p<0.001).

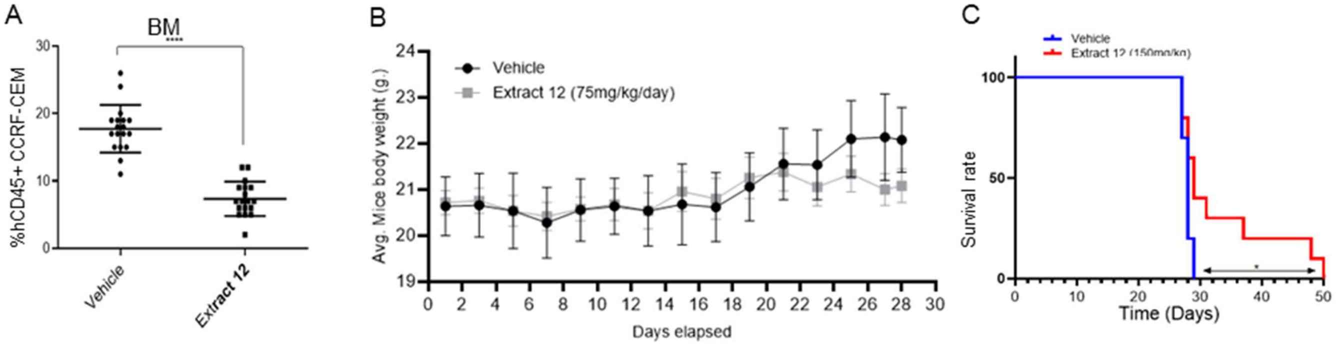


**fig. S6. Inhibition of tumor growth in-vivo by whole Extract.** (**A**) Female NSG mice were intravenously injected with $1\times10^6$ human CCRF-CEM cells. Mice were randomly divided into two groups (n=19) and alternate-day treated intraperitoneally with either vehicle or Extract 12 (150 mg/kg), after four weeks the percentage of hCD45-positive cells in the bone marrow (BM) was measured by flow cytometry and statistically analyzed by unpaired Student's *t*-test (****$p<0.0001$). (**B**) Body weight (grams) in vehicle- and extract-treated mice. (**C**) Survival analyses after CCRF-CEM injection followed by treatment with either vehicle or the whole extract (n=10/group), statistical differences were calculated with the Log-rank (Mantel-Cox) test (*$p<0.05$).

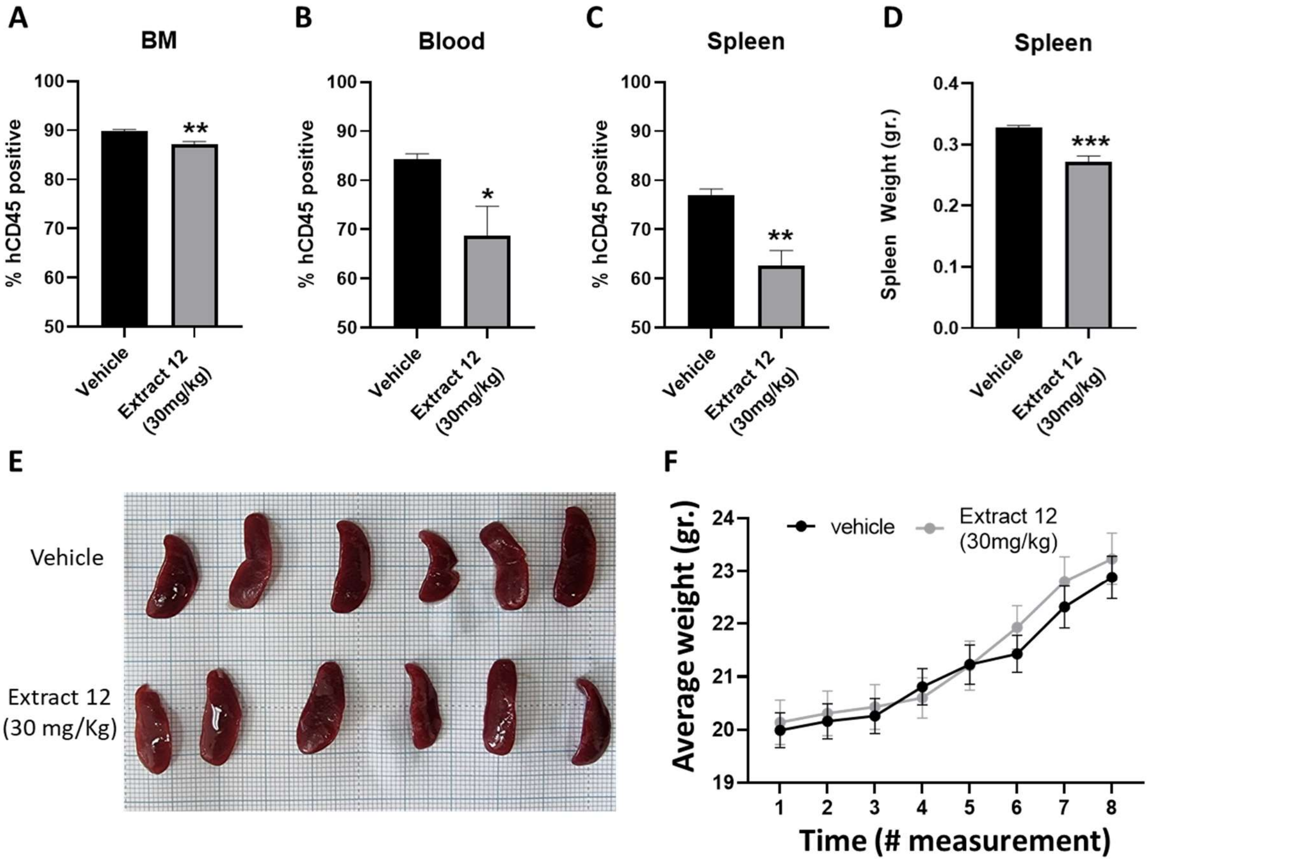


**fig. S7. Treatment with extract containing the three cannabinoids inhibits leukemic expansion in an *in-vivo* PDX model.** NSG mice were engrafted with $1\times10^6$ primary Notch1-mutated cells expanded from a T-ALL patient. After 35 days, the mice were randomly divided into two groups (N=6), a control group and an Extract 12-treatment group. The mice were treated on alternating days for 21 days and then evaluated for the percent of human CD45 positive cells from the (**A**) bone marrow, (**B**) peripheral blood and (**C**) spleen by flow cytometry. (D) The spleens were harvested and weighed. The average weight was compared between the two groups. (F) Representative image of the spleens from the vehicle treated (top) and Extract 12 treated (bottom) groups. (**F**) The mice were weighed every 5-8 days, starting on day 1. Data are presented as mean ± S.D. and statistically analyzed by a student's t-test (*p<0.05, **p<0.01, ***p<0.001).
